# Extracellular Matrix Abnormalities Contribute to Cardiac Insulin Resistance and Associated Dysfunction in Diet-induced Obese Mice

**DOI:** 10.1101/2023.11.14.567128

**Authors:** Vishal Musale, Colin E. Murdoch, Ayman K. Banah, Annie Hasib, Chandani K. Hennayake, Bo Dong, Chim C. Lang, David H. Wasserman, Li Kang

## Abstract

Increased deposition of extracellular matrix (ECM) components such as collagens and hyaluronan contributes to the pathogenesis of obesity-associated insulin resistance in muscle, liver, and adipose tissue. Despite the significance of the heart in cardiovascular and metabolic diseases, maladaptive ECM remodelling in obesity-associated cardiac insulin resistance and cardiac dysfunction has not been studied. Using genetic and pharmacological approaches in mice fed a high fat (HF) diet, we demonstrated a tight association between increased ECM deposition with cardiac insulin resistance. Increased collagen deposition by genetic deletion of matrix metalloproteinase 9 (MMP9) exacerbated cardiac insulin resistance and decreased hyaluronan deposition by treatment with PEGylated human recombinant hyaluronidase PH20 (PEGPH20) improved cardiac insulin resistance in obese mice. These relationships corresponded to functional changes in the heart. PEGPH20 treatment in obese mice ameliorated HF diet-induced abnormal myocardial remodelling. In addition to hyaluronan, increased collagen deposition is a characteristic of the obese mouse heart. We further demonstrated that pirfenidone, a clinically available anti-fibrotic medication which inhibits collagen expression, improved cardiac insulin resistance and cardiac function in obese mice. Our results provide important new insights into the role of ECM remodelling in the pathogenesis of cardiac insulin resistance and associated dysfunction in obesity of distinct mouse models. These findings support the novel therapeutic potential of targeting early cardiac ECM abnormalities in the prevention and treatment of obesity-related cardiovascular complications.

## Introduction

Insulin resistance associated with increasing prevalence of obesity, is defined as the inability of insulin to activate insulin signalling to effectively regulate multiple cellular processes including the promotion of glucose uptake and utilisation as fuels in the heart [1]. Importantly, cardiac insulin resistance contributes to myocardial dysfunction [2]. This is mediated by myocardial metabolic inflexibility, impaired calcium handling, mitochondrial dysfunction, dysregulated myocardial-endothelial interactions resulting in energy deficiency, impaired diastolic function, and myocardial cell death [3]. Although factors including oxidative stress, altered secretion of adipokines/cytokines, and neurohormonal activation in the renin-angiotensin system have been proposed to contribute to cardiac insulin resistance [1–3], the primary mechanisms underlying insulin resistance in the cardiovascular system are still not fully defined.

We have previously reported *in vivo* evidence that demonstrated increased expression of extracellular matrix (ECM) components contributed to the pathogenesis of obesity-associated insulin resistance in skeletal muscle [4, 5], liver [6], and adipose tissue [7]. Even though the heart is central to many obesity-associated disease states, maladaptive ECM remodelling in obesity-associated cardiac insulin resistance and related cardiac dysfunction has received little attention. The heart ECM comprises a complex network of macromolecules including proteins, proteoglycans, and growth factors, which provide structural and biochemical support to the surrounding cells and are essential for cellular and whole body homeostasis [8]. Myocardial fibrosis, characterised by excessive deposition of ECM components, leads to stiffening of the ventricles and negatively affects both contraction and relaxation of the heart [9]. Increased deposition of ECM components (e.g. collagens) has been shown to be key determinants of the increased left ventricular stiffness in patients with both heart failure with reduced ejection fraction and heart failure with preserved ejection fraction [10, 11]. It is noteworthy that remodelling of the heart ECM to a pro-fibrotic state can occur early and precede development of clinical fibrosis or cardiac histologic changes in hypertrophic cardiac conditions [11]. Therapies that can potentially inhibit the progression of these early changes of the ECM to severe fibrosis may also influence cardiac insulin resistance and preserve left ventricular function, thereby preventing the development of further cardiovascular complications.

In the present study the hypothesis that cardiac insulin resistance is associated with increased deposition of ECM components in the heart, leading to impaired cardiac function was tested. For this purpose, high fat (HF) diet fed, obese mouse models were studied. Further, we investigated whether pharmacological interventions that reduced heart ECM constituents using clinical and pre-clinical anti-fibrotic agents could reverse cardiac insulin resistance and improve cardiac function in obesity.

## Material and Methods

### Mouse models and treatment regimens

Animal experiments were carried out in compliance with the UK Animals (Scientific Procedures) Act 1986 and approved by the Animal Care and Use Committees of University of Dundee and Vanderbilt University. All the mice were maintained in an air-conditioned room (22 ± 2°C) with a 12:12-h light-dark cycle. Standard laboratory chow (13% calories as fat, LabDiet 5001) and tap water were supplied *ad libitum*, unless dietary or pharmacological interventions indicated otherwise.

#### High fat (HF) diet-induced obesity

Male C57BL/6J mice were purchased from the Jackson Laboratory. Following one week acclimatization period, mice starting from the age of 7 weeks were fed a high fat (HF) diet (60% calories as fat, SDS 824054 or BioServ F3282) for 16 weeks to induce obesity or maintained on chow diet and used as lean controls.

#### MMP9 knockout mice

The homozygous matrix metalloproteinases 9 (MMP9) null mice (*mmp9*^−/−^; Jackson Laboratory 007084) and their wild-type littermate controls (*mmp9*^+/+^) on a C57BL/6J background were fed a HF diet (60% calories as fat, BioServ F3282) starting at 3 weeks of age for 16 weeks. All mice were studied at 19 weeks of age. Cardiac insulin sensitivity was measured by hyperinsulinaemic-euglycaemic clamps [12].

#### PEGylated human recombinant hyaluronidase PH20 (PEGPH20) treatment

To investigate the cardiometabolic regulation of antifibrotic agents in obesity, after 12 weeks of HF diet feeding (60% calories as fat, SDS 824054) mice were treated with either vehicle (10 mmol/L histidine, 130 mmol/L NaCl at pH 6.5) or PEGPH20 (Provided under a Material Transfer Agreement with Halozyme Therapeutics, San Diego, CA) at 1 mg/kg through tail vein injections, once every 3 days for 24 days [13]. This PEGPH20 dose and treatment regimen have been previously shown to reduce hyaluronan in mice fed a HF diet to the levels seen in chow-fed lean mice. A PEGylated form of human recombinant PH20 was used due to its increased half-life [14]. Animals were maintained on HF diet during the treatment. Body weight was monitored at 3-day intervals. At the end of treatment, body composition was determined using EchoMRI (Echo Medical Systems, TX), cardiac insulin sensitivity was determined by hyperinsulinaemic-euglycaemic clamp [5], and left ventricular cardiac dynamics was determined by Pressure-Volume (PV) loop analysis (Transonic) using PV conductance catheter in closed chest preparation.

#### Pirfenidone Intervention

After 12 weeks of HF diet feeding (60% calories as fat, SDS 824054), mice received twice-daily treatments of either vehicle (0.25% carboxymethyl cellulose) or pirfenidone (125 mg/kg body weight) by oral gavage for 21 days, while animals were maintained on the HF diet. After the treatment, insulin sensitivity was measured by hyperinsulinaemic-euglycaemic clamp or cardiac function was analysed by PV loop in separate subgroups.

### Hyperinsulinaemic-euglycaemic clamp

Five to seven days prior to clamps, catheters were implanted in the carotid artery and jugular vein of mice for sampling and intravenous infusion, with the exception of mice used in the pirfenidone study, a jugular catheter was implanted for intravenous infusion and blood sampling was obtained from tail vein bleeding. Mice received constant infusion of insulin (4mU·kg^-1^·min^-1^) to achieve hyperinsulinaemia. Euglycaemia was maintained by assessing blood glucose every 10min and adjusting the glucose infusion rate (GIR) accordingly. Blood samples were taken at 80, 90, 100, 110, and 120min for the measurement of glucose rates of appearance (Ra) and disappearance (Rd) using a primed-infusion of [3-^3^H]glucose. At t=100 and 120min, plasma insulin concentrations were measured. At 120min, [^14^C]2-deoxyglucose intravenous bolus was administered and blood samples were collected from 2 to 35min after injection for the measurement of glucose uptake in tissues including the left-ventricle of the heart. Mice were euthanised after the last blood sample and tissues were excised for analysis of tissue [^14^C]2-deoxyglucose-phosphate, immunohistochemistry, Western blotting, and qRT-PCR.

#### Non-steady state calculation of glucose flux

Ra and Rd were calculated using non-steady-state equations [15]. The glucose metabolic index (Rg) was calculated for the measurement of tissue-specific glucose uptake as previously described [16].

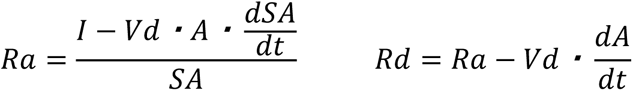

*Ra, Rd:* glucose appearance and disappearance rates (mg·kg^-1^·min^-1^); *I*: tracer infusion rate (dpm/min); *Vd*: volume distribution of glucose; *A*: concentration of glucose (mg/dL); *SA*: specific activity of glucose (dpm/mg); *t*: time (min). Endogenous glucose appearance rate (EndoRa) was calculated by subtracting the GIR from total Ra.

#### Glucose uptake (Rg) in tissues including the left ventricle of the heart was calculated as follows

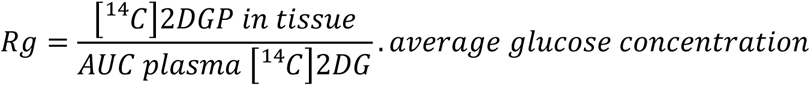

*[^14^C]-2DGP*: ^14^C-2-Deoxyglucose phosphate; *[^14^C]2DG*: 2-Deoxyglucose; *AUC:* Area under the curve.

Plasma insulin concentration was measured by Ultra-Sensitive Rat Insulin ELISA Kit (90060, CrystalChem). Plasma non-esterified fatty acid (NEFA) levels were measured by colorimetric analysis (434-91795, 436-91995, 4270-77000, WAKO Diagnostics).

### Pressure-Volume (PV) loop analysis

The real-time cardiac function of experimental mice was evaluated by left ventricle PV loops using an admittance catheter (1.2F, Transonic) coupled to ADV500 data acquisition system (Transonic) visualised by LabChart (ADInstruments). Mice were anaesthetized with 2% isoflurane (volume/volume) and the body temperature of mice was monitored by a rectal thermometer probe. The PV catheter equipped with both pressure and volume sensors was introduced into the aorta via the carotid artery to measure arterial pressure. The catheter was then advanced to the left ventricle to record pressure and volume signals, under basal and inferior vena cava (IVC) occlusion conditions as described previously [17]. IVC occlusion, a gold standard for load-independent measurements of contractility (end-systolic pressure-volume relationship (ESPVR)) and compliance (end-diastolic pressure-volume relationship (EDPVR)) was performed by obstructing the return flow of blood to the heart. The load-dependent and independent hemodynamic data obtained from the experimental mice were analysed using Lab Chart Pro 8 software (ADInstruments).

### Immunohistochemistry

Paraffin-embedded left ventricular tissues were cut into a 5-6µm section using a microtome and mounted onto the staining slide. Hyaluronan, collagen, CD31, CD45, and α-SMA expressions were assessed using biotinylated hyaluronan-binding protein (AMS.HKD-BC41, AMS Biotechnology), Picrosirius Red (Direct Red 80, Sigma 365548), anti-CD31 antibody (NBP1-49805, Novus Biologicals), anti-CD45 antibody (BD Bioscience 550539), or anti-α-SMA antibody (Cell Signalling 19245), respectively. Cardiomyocyte area and capillaries were co-stained with wheat-germ agglutinin (WGA, 2BSCIENTIFIC RL-1022-5) and isolectin B4 (Vector B-1205, 2BSCIENTIFIC B-1205-05). For the staining of hyaluronan, CD31, CD45 and α-SMA, slides were lightly counterstained with Mayer’s hematoxylin. The specificity of hyaluronan staining was confirmed by treating the sample section with or without recombinant human hyaluronidase PH20. For CD31 staining, bowel samples were used as a positive control. Images (10-12 images per animal) were captured by Axiovision microscope (Zeiss Axioscope, Germany) and quantified using Image J software. Hyaluronan and collagen content was measured as the percentage of total left ventricular area under polarized light. Capillary density was quantified as the number of capillaries (CD31-positive structures) per square millimetres. CD45 and α-SMA positive cells were counted as cells per square millimetres.

### Western Blotting

Left ventricles of mouse hearts were dissected and homogenized in lysis buffer containing protease and phosphatase inhibitors as previously described [7]. Protein concentrations were determined and 20-40µg of protein per sample was loaded onto 4-12% SDS-PAGE gels for protein detection, using antibodies against CD44 (AF6127, 1:1,000; R&D Systems), RHAMM (87129, 1:1000, Cell Signalling), TGF-β (ab179695, 1:1000, Abcam), Phospho-Smad2 (Ser465/467)/Smad3 (Ser423/425) (8828, 1:1000, Cell signalling), Smad2/3 (3102, 1:1000, Cell signalling), VCAM-1 (AF643, 1:1000, R&D Systems), BNP (ab19645, 1:1000, Abcam), Phospho-p38 MAPK (Thr180/Tyr182) (9211, 1:1000, Cell signalling), p38 MAPK (9212, 1:1000, Cell signalling), Phospho-SAPK/JNK (Thr183/Tyr185) (9251, 1:1000, Cell signalling), JNK (9252, 1:1000, Cell signalling), pERK1/2 (4370, 1:1000, Cell signalling), ERK (4695, 1:1000, Cell signalling), α-SMA (19245, 1:1000, Cell signalling), pAKT (9271, 1:1000, Cell signalling), and AKT (9272, 1:1000, Cell signalling). GAPDH (5174, 1:1000, Cell signalling), beta-tubulin (ab6046, 1:1000, Abcam), and ponceau staining were used as loading controls.

### Quantitative real-time PCR

Total RNA was extracted using TriPure isolation reagent and reversed transcribed into cDNA using SuperScript™ II Reverse Transcriptase (18064014, ThermoFisher). Quantitative real-time PCR was carried out to amplify genes of interest using the Veriti 96-well Thermal Cycler, ThermoFisher. Primer sequences can be found in Supplemental Table 1. Data were normalized to 18S gene expression and analysed using the 2^-ΔΔCT^ method.

### Statistical analysis

Data are presented as mean ± S.E.M. Statistical analysis was performed using unpaired Student t-test or either one-way or two-way ANOVA followed by the Tukey’s method for multiple comparisons where appropriate. The significance level was set at p<0.05.

## Results

### Increased ECM deposition in the heart is associated with cardiac insulin resistance in obesity

Protein expression of collagen III and collagen IV, and hyaluronan content in the heart were increased (or tended to be increased) in HF-fed obese mice when compared to chow-fed mice (Fig 1A-D). HF diet feeding in mice induced cardiac insulin resistance as shown by decreased glucose uptake in the heart during an hyperinsulinaemic-euglycaemic clamp (insulin clamp) (Fig 1E) [13]. Genetic deletion of MMP9 increased protein expression of collagen III and tended to increase protein expression of collagen IV in HF-fed mice when compared with HF-fed wildtype controls (Fig 1F-H). Increased collagen deposition in the HF-fed *MMP9*^-/-^ mice was accompanied by exacerbated cardiac insulin resistance with decreased glucose uptake in heart during an insulin clamp (Fig 1I) [12]. In contrast, pharmacological treatment of PEGPH20 decreased hyaluronan content in the heart of HF-fed obese mice which was accompanied by improved cardiac insulin resistance as evidenced by increased cardiac glucose uptake during an insulin clamp (Fig 1J-L) [5]. These results suggest that obesity led to an increased ECM deposition in the heart which was tightly associated with cardiac insulin resistance in mice.

**Figure 1.**
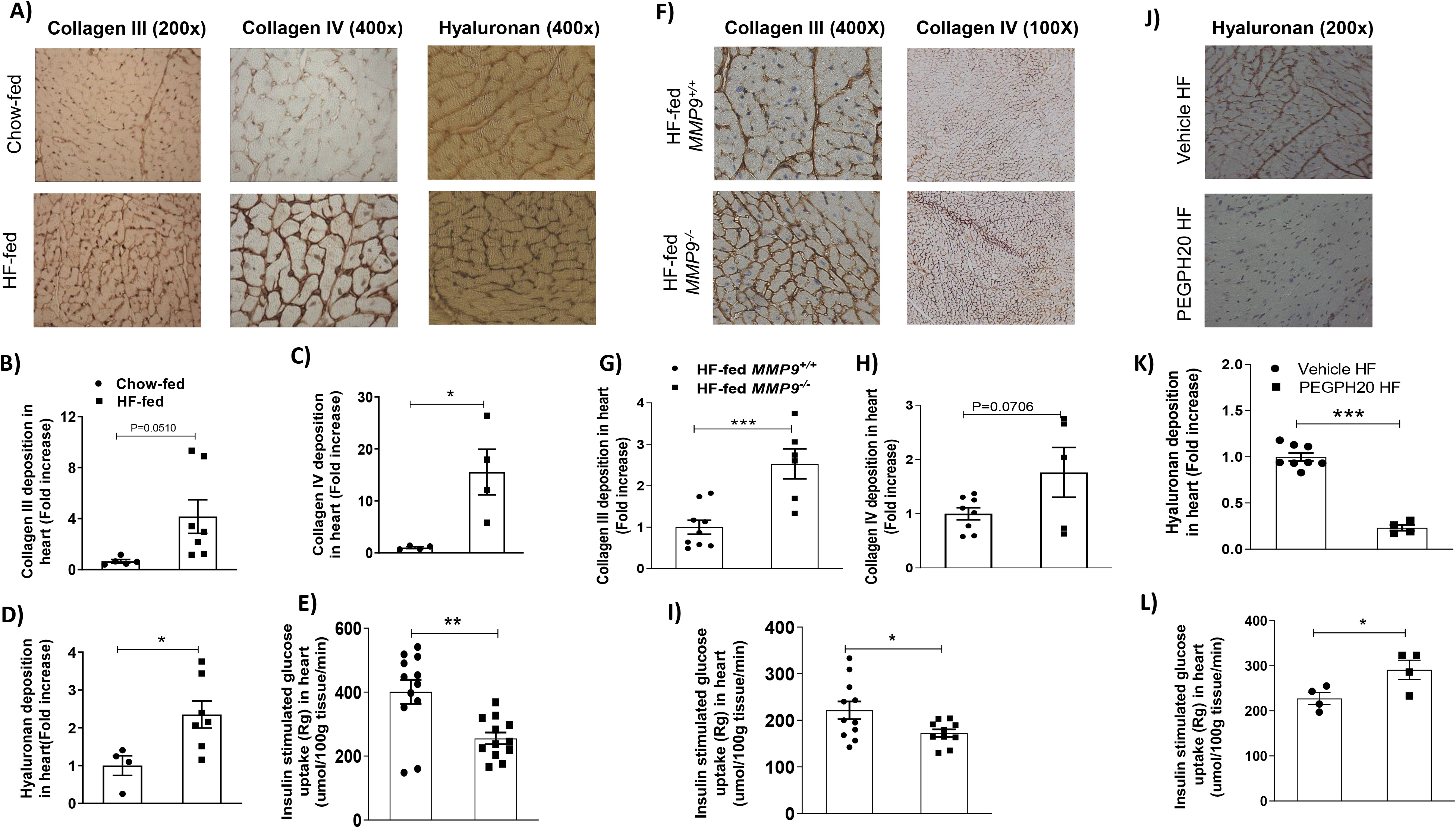
Increased deposition of ECM collagen and hyaluronan was associated with cardiac insulin resistance in obese mice. (A-D) C57BL/6 mice were fed either a chow diet or a 60% high fat (HF) diet for 16 weeks. Collagen III, collagen IV, and hyaluronan were detected by immunohistochemistry and quantified by ImageJ in heart sections. Representative images were shown. N=4-5 for chow-fed mice, and n=4-7 for HF-fed mice. (E) Cardiac insulin sensitivity was assessed by insulin-stimulated glucose uptake during a hyperinsulinemic-euglycemic clamp (n=12). (F-H) The homozygous MMP9 knockout mice (*MMP9*^-/-^) and their wildtype littermate controls (*MMP9*^+/+^) were fed with 60% HF diet for 16 weeks. Collagens III and IV were detected by immunohistochemistry and quantified by ImageJ in heart sections. Representative images were shown (1.55 pixels/μm for 100x images; 3.10 pixels/μm for 200x images; 6.21 pixels/μm for 400x images). N=8-9 for HF-fed *MMP9*^+/+^, and n=5-6 for HF-fed *MMP9*^-/-^. (I) Cardiac insulin sensitivity was assessed by insulin-stimulated glucose uptake during a hyperinsulinemic-euglycemic clamp (n=11). (J) C57BL/6 mice were fed a 60% HF diet for 12 weeks before receiving either vehicle or PEGPH20, once every 3 days for 24 days. Hyaluronan was detected by immunohistochemistry and quantified by ImageJ in heart sections. Representative images were shown. N=8 for Vehicle HF and n=4 for PEGPH20 HF. (L) Cardiac insulin sensitivity was assessed by insulin-stimulated glucose uptake during a hyperinsulinemic-euglycemic clamp (n=4).\**p*<0.05, \*\**p*<0.01, and \*\*\**p*<0.005.

### Reduction of hyaluronan ameliorates obesity-associated cardiac dysfunction

To determine whether the association between increased ECM deposition and cardiac insulin resistance also extends to cardiac dysfunction in obesity, *in vivo* cardiac performance was measured in a separate group of mice fed with HF diet and treated with PEGPH20. PEGPH20 treatment decreased total body mass and %fat mass but increased %lean mass in HF-fed obese mice (Supplemental Fig 1A-B). Hyaluronan content in the heart was decreased by PEGPH20 (Supplemental Fig 1C-D).

HF diet feeding in mice led to cardiac dysfunction (Table 1). Systolic, diastolic and pulse pressures were increased in HF-fed vehicle-treated (HF-Vehicle) mice when compared with lean control mice. End systolic pressure (Pes) was significantly elevated, and there was a tendency albeit not significant (p=0.07) for elevation of end diastolic pressure (Ped) after HF diet. Interestingly, the HF diet induced an inotropic effect in the left ventricle, with significantly higher dP/dt (max and min) and a tendency for an increase in a load-independent mesurement of contractility (end systolic pressure-volume relationship (ESPVR), p=0.07). The arterial elastance (Ea) was also increased in HF-Vehicle mice in comparison with the lean controls, implying an impaired ventricular arterial coupling. Taken together, mice fed a HF diet underwent abnormal myocardial remodelling, working under higher pressures (Pes) with an increased afterload (Ea) and an increased inotropic response (dP/dt and ESPVR). Diastolic function was not altered by HF diet with the relaxation constant (Tau) and end diastolic pressure-volume relationship (EDPVR) remaining similar between HF-Vehicle and lean control mice.

**Table 1.** Hemodynamic parameters from the pressure-volume loop analyses of chow-fed lean control mice and high fat (HF)-fed mice receiving vehicle or PEGPH20. N=7 for Lean-Control, n=10 for HF-Vehicle, and n=11 for HF-PEGPH20. \**p*<0.05, \*\**p*<0.01, \*\*\**p*<0.01 compared with Lean-Control; ^Δ^*p*<0.05, ^ΔΔ^*p*<0.01 compared with HF-Vehicle; ^#^*p*=0.07 compared with Lean-Control. IVC: inferior vena cava occlusion; ESPVR: end systolic pressure volume relationship; EDPVR: end diastolic pressure volume relationship; VAC: ventricular-arterial coupling; LV: left ventricle.

| Parameters | Lean-Control | HF-Vehicle | HF-PEGPH20 |
| --- | --- | --- | --- |
| **Blood Pressures** |  |  |  |
| Systolic blood pressure (mmHg) | 97.04 ± 2.68 | 112.4 ± 3.53 ** | 98.45 ± 2.81 $^{\Delta\Delta}$ |
| Diastolic blood pressure (mmHg) | 68.12 ± 1.19 | 77.70 ± 2.26 ** | 67.06 ± 2.63 $^{\Delta\Delta}$ |
| Pulse pressure (mmHg) | 30.27 ± 0.59 | 37.35 ± 2.15 ** | 31.57 ± 0.38 $^{\Delta\Delta}$ |
| **Baseline Parameters** |  |  |  |
| Heart rate | 554.7 ± 7.81 | 545.5 ± 8.43 | 559.9 ± 11.37 |
| End systolic volume (Ves) (μL) | 16.68 ± 1.80 | 13.38 ± 2.30 | 17.63 ± 3.60 |
| End diastolic volume (Ved) (μL) | 29.41 ± 2.31 | 23.46 ± 2.49 | 32.36 ± 2.33 $^{\Delta}$ |
| End systolic pressure (Pes) (mmHg) | 92.53 ± 2.69 | 113.4 ± 3.89 *** | 94.74 ± 2.97 $^{\Delta\Delta}$ |
| End diastolic pressure (Ped) (mmHg) | 6.718 ± 1.08 | 10.90 ± 1.35 # | 6.489 ± 1.22 $^{\Delta}$ |
| Stroke volume (SV) (μL) | 22.03 ± 1.63 | 16.25 ± 1.46 | 23.59 ± 2.51 $^{\Delta}$ |
| Ejection Fraction (EF) (%) | 60.91 ± 3.23 | 55.23 ± 3.82 | 62.08 ± 4.54 |
| Cardiac output (CO) (μL/min) | 12250 ± 890.9 | 8799 ± 713.6 | 13170 ± 1392 $^{\Delta}$ |
| Stroke work (SW) (mmHg*μL) | 1953 ± 130.9 | 1644 ± 131.5 | 2136 ± 222.6 |
| dP/dt max (mmHg/s) | 8196 ± 580.1 | 10470 ± 348.4 * | 8895 ± 224.5 $^{\Delta}$ |
| dP/dt min (mmHg/s) | 8083 ± 400.5 | 10360 ± 650.7 ** | 8045 ± 258.8 $^{\Delta\Delta}$ |
| Tau (ms) | 5.794 ± 0.31 | 6.528 ± 0.36 | 5.961 ± 0.34 |
| **IVC** |  |  |  |
| ESPVR | 4.138 ± 0.31 | 5.746 ± 0.44 # | 4.158 ± 0.44 |
| EDPVR | 0.06494 ± 0.01 | 0.06211 ± 0.01 | 0.05274 ± 0.01 |
| **VAC** |  |  |  |
| Arterial elastance (Ea) | 4.286 ± 0.39 | 6.303 ± 0.45 * | 4.502 ± 0.45 $^{\Delta}$ |
| End systolic elastance (Ees) | 5.385 ± 0.53 | 8.977 ± 1.26 | 7.186 ± 1.04 |
| VAC index | 1.081± 0.18 | 0.9814± 0.14 | 0.6301± 0.07 |
| **Clinical Parameters** |  |  |  |
| Heart weight/Femur (mg/mm) | 9.76 ± 0.85 | 11.16 ± 0.64 | 10.01 ± 0.37 |
| LV/Femur (mg/mm) | 7.83 ± 0.74 | 8.02 ± 0.55 | 7.83 ± 0.29 |

Mice fed a HF diet for 12 weeks were administered PEGPH20 for 24 days while remaining on HF diet. PEGPH20 intervention prevented diet-induced myocardial remodelling (Table 1). HF diet-induced increases in systolic, diastolic, and pulse pressures were eliminated by PEGPH20 treatment. PEGPH20 decreased Pes and Ped, increased end diastolic volume (Ved), stroke volume (SW) and cardiac output (CO). PEGPH20 corrected the HF diet-induced changes in dP/dt max, dP/dt min, ESPVR, and Ea. Other parameters such as heart rate, ejection fraction, stroke work, Tau, end systolic elastance (Ees), EDPVR, and ventricular arterial coupling index were not significantly different between groups. Neither HF diet feeding nor PEGPH20 treatment changed whole heart or left ventricle weights. These results suggest that PEGPH20 treatment prevented abnormal myocardial remodelling observed in mice after HF diet feeding.

### Removal of hyaluronan reduces HF diet-induced cardiac hypertrophy, SMAD activation, and inflammation

HF diet feeding in mice increased both interstitial and perivascular collagen deposition in the left ventricle (Fig 2A-B). These increases were absent in PEGPH20-treated HF-fed (HF-PEGPH20) mice. However, protein expression of α-SMA (smooth muscle actin), a marker of cardio-myofibroblast activation, was not different between mouse groups (Fig 2C). Concurrently, HF diet feeding in mice increased the cross-sectional area of cardiomyocytes, indicative of hypertrophy. This effect was partially restored by PEGPH20 treatment (Fig 2D-E). We also observed a notable decrease in both capillary density and the number of cardiomyocytes per mm^2^ in HF-Vehicle mice compared with lean controls (Fig 2D, F-G). These decreases were not significantly changed by PEGPH20.

**Figure 2.**
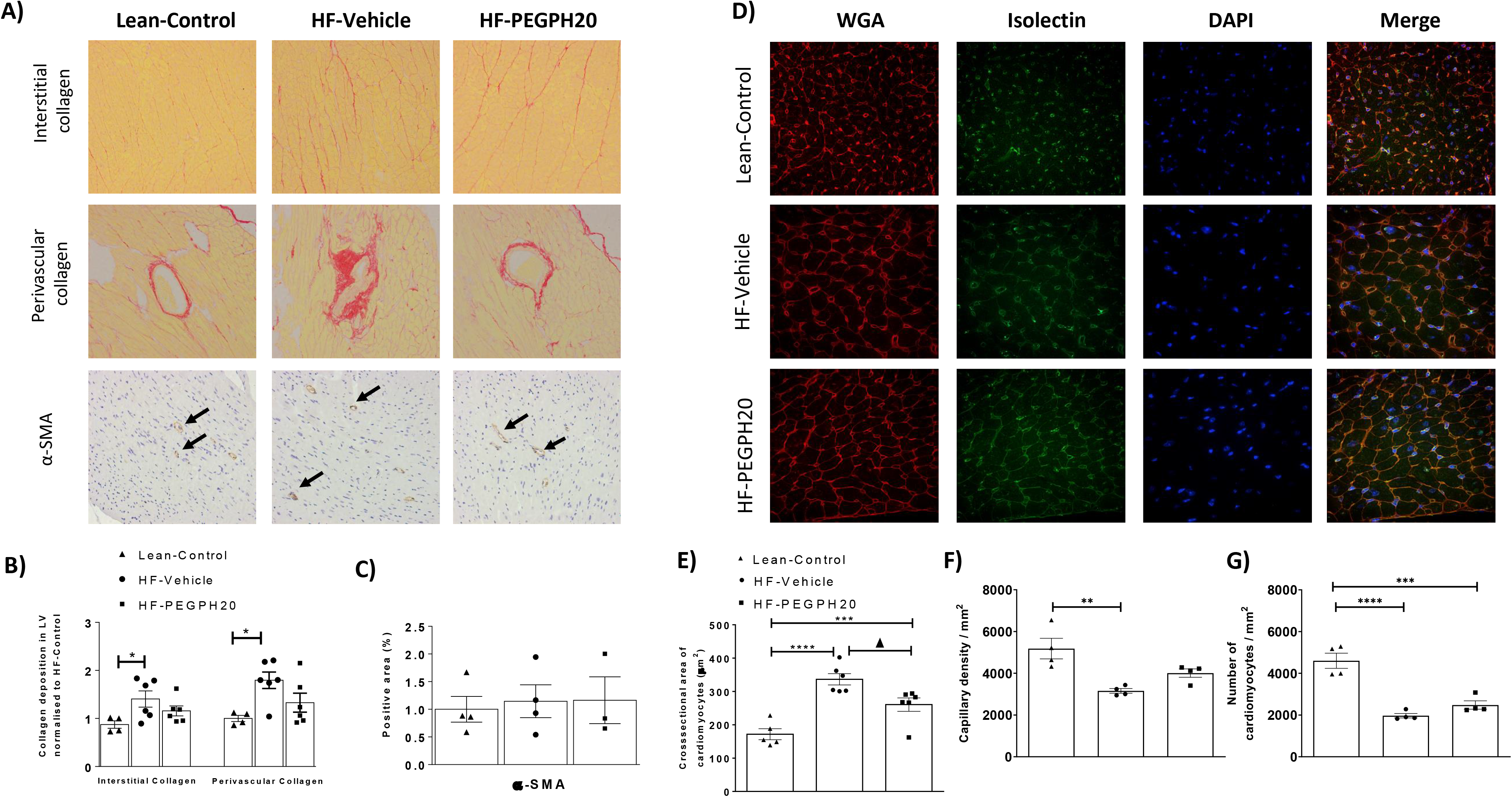
PEGPH20 treatment ameliorated high fat (HF) diet-induced myocardial fibrosis and hypertrophy. C57BL/6 mice were fed either a chow diet or a 60% HF diet for 16 weeks. After 12 weeks of HF feeding, HF-fed mice received either vehicle or PEGPH20, once every 3 days for 24 days. (A-C) Interstitial and perivascular collagens were detected by Sirius Red staining in left ventricle sections. Expression of α-SMA (smooth muscle actin) was detected by immunohistochemistry. Data were quantified by ImageJ. N=4-6. (D-G) Cardiomyocyte size was determined by Wheat Germ Agglutinin (WGA) staining. Capillary density was assessed by isolectin staining. DAPI was used to stain cell nuclei. Representative images were shown at 200x magnification (3.10 pixels/μm). Images were quantified by ImageJ. N=4-6. \**p*<0.05, \*\**p*<0.01, \*\*\**p*<0.005, and \*\*\*\**p*<0.001 compared with Lean-Control; ^Δ^*p*<0.05 compared with HF-Vehicle.

We further studied cellular signalling changes in the left ventricle associated with HF feeding and PEGPH20 treatment. Protein expression of CD44, one of the two primary hyaluronan receptors, was decreased in HF-Vehicle mice compared to lean controls. CD44 remained low in HF-PEGPH20 mice (Fig 3A-B). Protein expression of RHAMM, the other hyaluronan receptor was not changed in HF-Vehicle mice but was increased in HF-PEGPH20 mice relative to lean controls (Fig 3A, C). While TGF-β, total SMAD2/3, and phosphorylated SMAD2/3 were not different between groups, the ratio of pSMAD2/3 to total SMAD2/3 was significantly decreased in HF-PEGPH20 mice relative to HF-Vehicle mice, suggesting a decreased SMAD2/3 activation (Fig 3A, D-G). VCAM-1, vascular cell adhesion molecule 1, a marker of inflammation-associated vascular adhesion was increased by HF diet feeding. This effect of HF diet was reversed by PEGPH20 treatment (Fig 3A, H). BNP, brain natriuretic peptide, a biomarker of cardiac function was not different between groups (Fig 3A, I). MAPK signalling was not altered by HF feeding or PEGPH20 treatment independently. However, phosphorylated ERK1/2 and the ratio of pERK1/2 to total ERK1/2 were lower in HF-PEGPH20 mice relative to lean control mice (Supplemental Fig 2).

**Figure 3.**
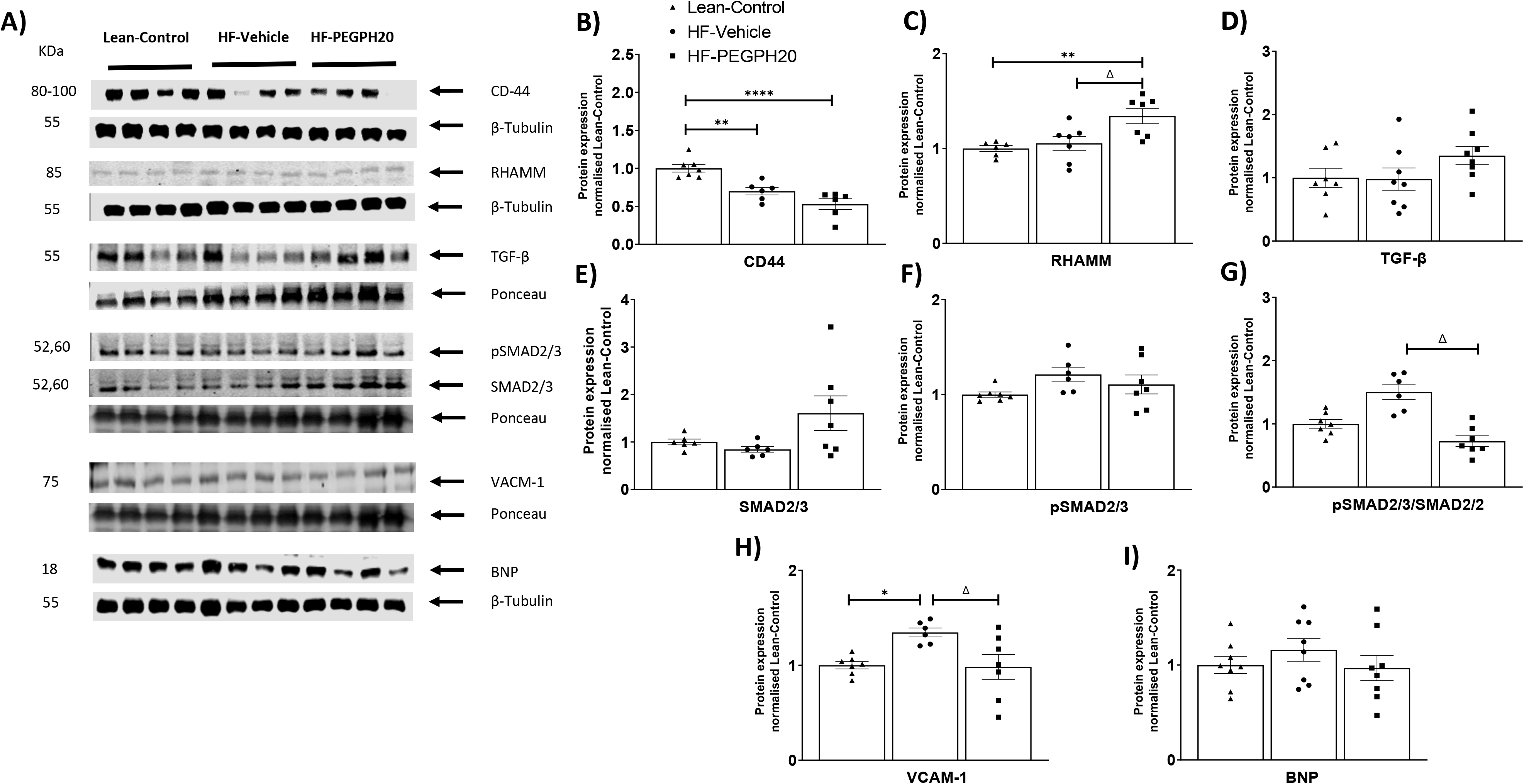
PEGPH20 treatment reduced SMAD2/3 activation and inflammation in the left ventricle of obese mice. C57BL/6 mice were fed either a chow diet or a 60% HF diet for 16 weeks. After 12 weeks of HF feeding, HF-fed mice received either vehicle or PEGPH20, once every 3 days for 24 days. Protein expression was determined by Western blotting. Representative blots were shown. N=6-8. \**p*<0.05, \*\**p*<0.01, \*\*\**p*<0.005, and \*\*\*\**p*<0.001 compared with Lean-Control; ^Δ^*p*<0.05 compared with HF-Vehicle. BNP: brain natriuretic peptide.

HF diet feeding in mice increased cardiac inflammation as evidenced by increased gene expression of TNF-α and increased CD45^+^ cells in the left ventricle. The increase in inflammatory markers was prevented by PEGPH20 treatment (Fig 4A, G-H). mRNA levels of IL-1β, IL-6, IL-10, BNP and β-MHC were not significantly different between groups (Fig 4B-F).

**Figure 4.**
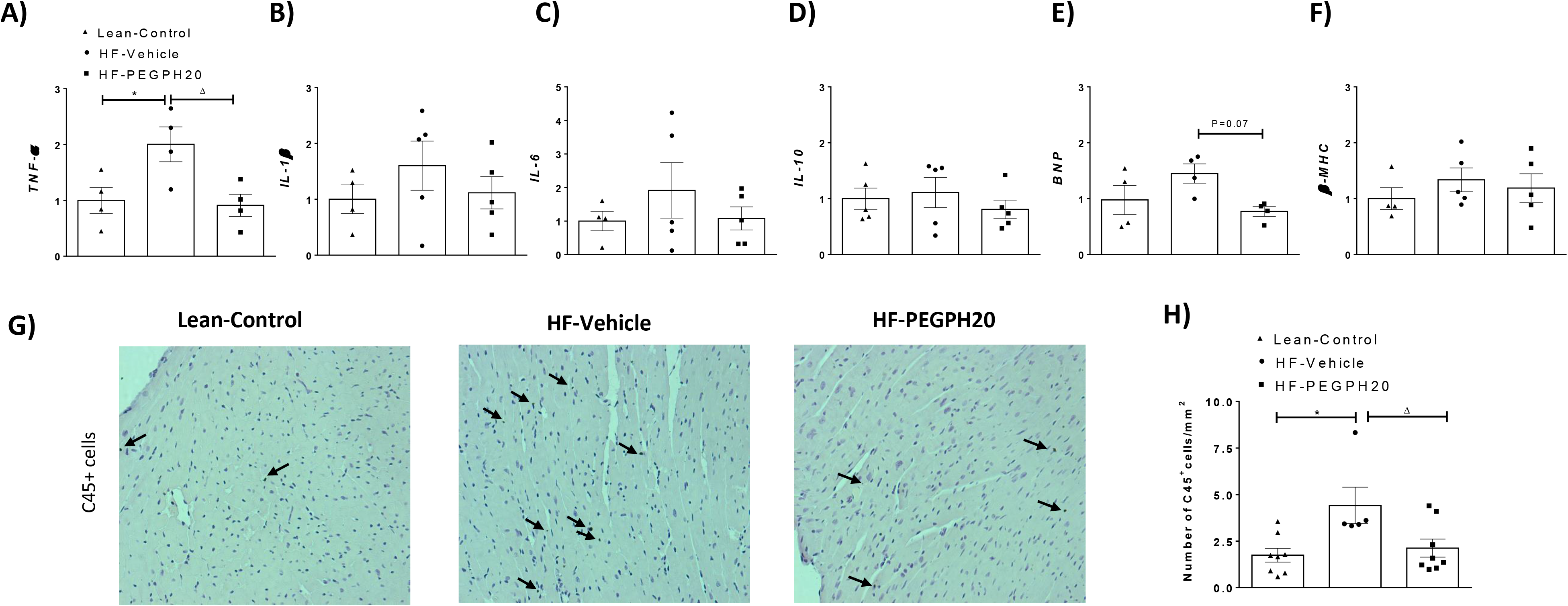
PEGPH20 treatment reduced high fat (HF) diet-induced inflammation in the left ventricle of obese mice. C57BL/6 mice were fed either a chow diet or a 60% HF diet for 16 weeks. After 12 weeks of HF feeding, HF-fed mice received either vehicle or PEGPH20, once every 3 days for 24 days. (A-F) mRNA expression was determined by qRT-PCR. N=4-5. (G-H) CD45 positive cells were determined by CD45 immunohistochemistry and quantified by ImageJ. Representative images were shown at 200x magnification (3.10 pixels/μm). N=5-8. \**p*<0.05 compared with Lean-Control; ^Δ^*p*<0.05 compared with HF-Vehicle. BNP: brain natriuretic peptide; β-MHC: myosin heavy chain β.

### Pirfenidone ameliorates cardiac insulin resistance in obese mice

In light of the increased collagen deposition in HF-fed mice, we tested whether pirfenidone, an anti-fibrotic drug which has been shown to decrease collagen deposition in the heart [18–21], has beneficial effects on cardiac insulin resistance and associated cardiac dysfunction in obese mice. Pirfenidone treatment did not change body weight or body composition of HF-fed obese mice (Fig 5A-B). During oral glucose tolerance tests, pirfenidone-treated HF-fed mice exhibited a modest improvement in glycaemic response compared to vehicle-treated HF-fed mice (Fig 5C). However, there was no difference in area under the glucose curve (AUC) or plasma insulin levels between the two groups (Fig 5D-E). During insulin clamps, the arterial glucose levels were clamped at 6.5 mmol/L at the steady state of the clamps (80-120 min) in all the mice (Fig 5F). Pirfenidone treatment increased glucose infusion rates (Fig 5G), but the clamp insulin was significantly lower in pirfenidone-treated mice than in vehicle-treated HF mice, indicating an improvement in insulin action in pirfenidone-treated mice (Fig 5H). Insulin increased the rate of glucose disappearance (Rd) and decreased the rate of endogenous glucose appearance (EndoRa) in all mice, but to a much greater extent in pirfenidone-treated mice compared to vehicle-treated mice (Fig 5I-J). In addition, Rg, a measure of tissue-specific glucose uptake was significantly higher in left ventricle, vastus lateralis muscle, and subcutaneous white adipose tissue of pirfenidone-treated mice than vehicle-treated mice (Fig 5K). Taken together, these results suggest that pirfenidone treatment improved systemic as well as cardiac insulin resistance in obese mice.

**Figure 5.**
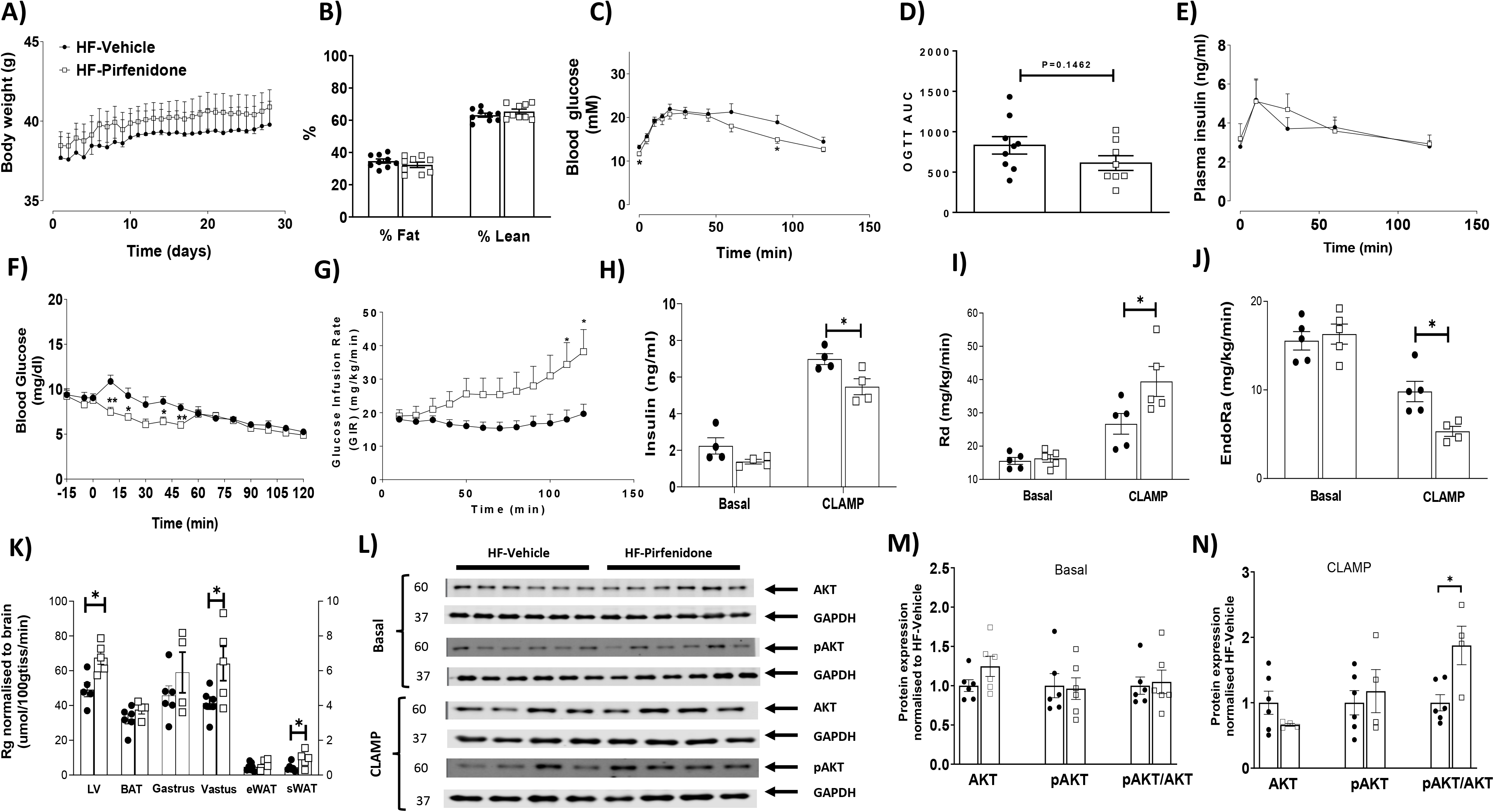
Pirfenidone improved cardiac as well as systemic insulin resistance in obese mice. C57BL/6 mice were fed with a 60% high fat (HF) diet for 16 weeks. After 12 weeks of HF feeding, mice received twice-daily treatments of vehicle or pirfenidone for 21 days. (A-B) Body weight was monitored daily and body composition was measured after the vehicle/drug treatment. N=9-10. (C-E) Blood glucose, area under the curve of blood glucose, and plasma insulin were determined during an oral glucose tolerance test. N=8-10. (F-K) Insulin sensitivity was determined by hyperinsulinemic-euglycemic clamps. (F) Blood glucose and (G) glucose infusion rate (GIR) were measured. N=5-6. (H) Plasma insulin concentrations were measured at basal state as well as during the insulin clamp. N=4. (I-J) Glucose disappearance (Rd) and endogenous glucose appearance (EndoRa) rates were measured during the clamp. N=4-5. (K) Tissue-specific glucose uptake (Rg) was measured during the clamp N=4-6. (L-N) Insulin signalling in the left ventricle was measured by pAKT/AKT ratio at both basal condition as well as post the insulin clamp, by Western blotting. Representative blots were shown and quantified. N=4-6. \**p*<0.05.

Consistent with increased cardiac insulin action, insulin signalling as determined as the ratio of pAKT/AKT was not changed at the basal state but was increased during insulin stimulation in the left ventricle of pirfenidone-treated mice relative to vehicle-treated mice (Fig 5M-N).

### Pirfenidone modestly prevents HF diet-induced myocardial remodelling in obese mice, in association with decreases in collagen deposition, SMAD activation, MAPK activation, and cardiac inflammation

We next studied whether improved cardiac insulin resistance by pirfenidone was associated with reversal of obesity-associated abnormal myocardial remodelling. Pirfenidone decreased pulse pressure and dP/dt min, both of which were previously shown to be increased by HF diet in mice (Table 2). However, no other parameters were significantly changed by pirfenidone (Table 2). Heart weight and left ventricular weight also remained the same between pirfenidone and vehicle-treated mice (Table 2).

**Table 2.** Hemodynamic parameters from the pressure-volume loop analyses of high fat (HF)-fed mice receiving vehicle or pirfenidone. N=9 for HF-Vehicle, and n=8 for HF-Pirfenidone. \*\**p*<0.01, \*\*\**p*<0.005 compared with HF-Vehicle. IVC: inferior vena cava occlusion; ESPVR: end systolic pressure volume relationship; EDPVR: end diastolic pressure volume relationship; VAC: ventricular-arterial coupling; LV: left ventricle.

| Parameters | HF-Vehicle | HF-Pirfenidone |
| --- | --- | --- |
| **Blood Pressures** |  |  |
| Systolic blood pressure (mmHg) | 93.96 ± 2.64 | 91.87 ± 1.54 |
| Diastolic blood pressure (mmHg) | 60.53 ± 1.85 | 64.08 ± 1.34 |
| Pulse pressure (mmHg) | 32.76 ± 0.65 | 27.08 ± 0.97 *** |
| **Baseline Parameters** |  |  |
| Heart rate | 557.1 ± 8.89 | 568.0 ± 9.20 |
| End systolic volume (Ves) (µL) | 10.30 ± 1.88 | 15.14 ± 1.74 |
| End diastolic volume (Ved) (µL) | 28.14 ± 1.83 | 34.93 ± 3.48 |
| End systolic pressure (Pes) (mmHg) | 84.78 ± 4.28 | 88.76 ± 2.14 |
| End diastolic pressure (Ped) (mmHg) | 7.558 ± 0.79 | 8.070 ± 0.85 |
| Stroke volume (SV) (µL) | 20.43 ± 1.40 | 20.86 ± 1.99 |
| Ejection Fraction (EF) (%) | 71.90 ± 4.34 | 61.69 ± 2.16 |
| Cardiac output (CO) (µL/min) | 11410 ± 1053 | 11890 ± 1219 |
| Stroke work (SW) (mmHg*µL) | 1802 ± 103.5 | 1776 ± 166.2 |
| dP/dt max (mmHg/s) | 9887 ± 242.4 | 9515 ± 374.2 |
| dP/dt min (mmHg/s) | 9011 ± 216.4 | 7488 ± 463.4 ** |
| Tau (ms) | 5.851 ± 0.26 | 6.197 ± 0.30 |
| **IVC** |  |  |
| ESPVR | 6.260 ± 0.93 | 5.468 ± 1.00 |
| EDPVR | 0.08802 ± 0.01 | 0.1091 ± 0.01 |
| **VAC** |  |  |
| Arterial elastance (Ea) | 4.419 ± 0.44 | 4.331 ± 0.48 |
| End systolic elastance (Ees) | 9.179 ± 1.90 | 6.707 ± 0.99 |
| VAC index | 0.4918 ± 0.09 | 0.6840 ± 0.06 |
| **Clinical Parameter** |  |  |
| Heart weight/Femur (mg/mm) | 10.71 ± 1.25 | 10.36 ± 0.48 |
| LV/Femur (mg/mm) | 7.32 ± 0.30 | 7.89 ± 0.24 |

Interestingly, pirfenidone decreased interstitial, but not perivascular collagen deposition (Fig 6A-B). Protein expression of α-SMA, the cross-sectional area of cardiomyocytes, capillary density, and the number of cardiomyocytes per mm^2^ were unchanged by pirfenidone (Fig 6C-G). Pirfenidone treatment decreased pSMAD2/3/SMAD2/3 ratio without affecting TGF-β expression (Fig 7A-C). Pirfenidone also did not affect α-SMA or VCAM-1 expression but decreased BNP expression by Western blotting (Fig 7A, D-F). In addition, pirfenidone treatment caused a significant decrease in total P38, a trend for a decrease in pP38, and a decrease in pJNK, without affecting ERK signalling (Fig 7A, G-I). Moreover, Pirfenidone decreased mRNA levels of TNF-α, IL-6, and BNP, without affecting mRNA levels of IL-1β, IL-10 and β-MHC, or numbers of CD45^+^ cells (Fig 8).

**Figure 6.**
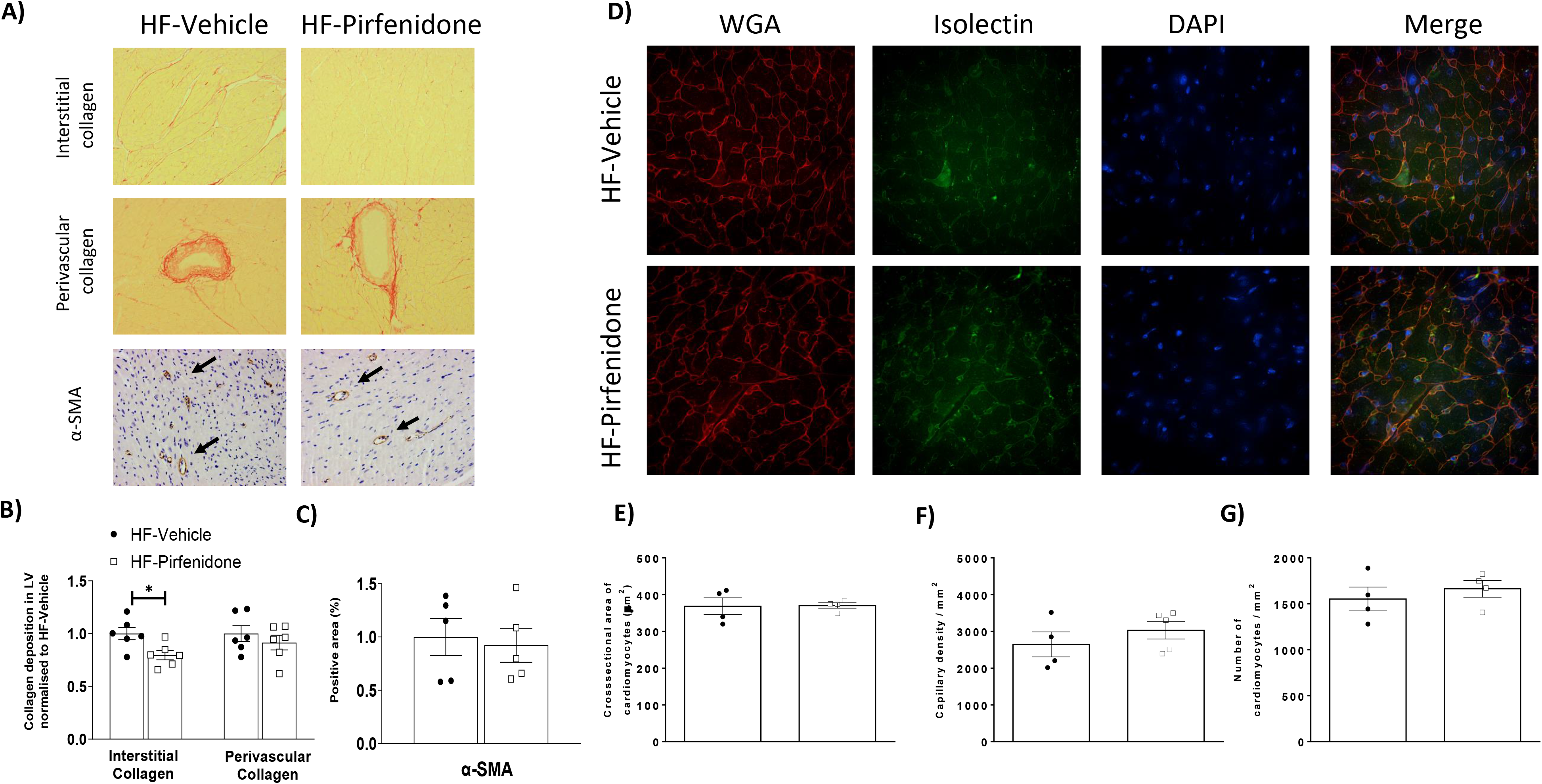
Pirfenidone reduced interstitial collagen deposition in the left ventricle of obese mice. C57BL/6 mice were fed with a 60% high fat (HF) diet for 16 weeks. After 12 weeks of HF feeding, mice received twice-daily treatments of vehicle or pirfenidone for 21 days. (A-C) Interstitial and perivascular collagens were detected by Sirius Red staining in left ventricle sections. Expression of α-SMA (smooth muscle actin) was detected by immunohistochemistry. Data were quantified by ImageJ. N=5-6. (D-G) Cardiomyocyte size was determined by Wheat Germ Agglutinin (WGA) staining. Capillary density was assessed by isolectin staining. DAPI was used to stain cell nuclei. N=4-5. Representative images were shown at 200x magnification (3.10 pixels/μm). Images were quantified by ImageJ. \**p*<0.05.

**Figure 7.**
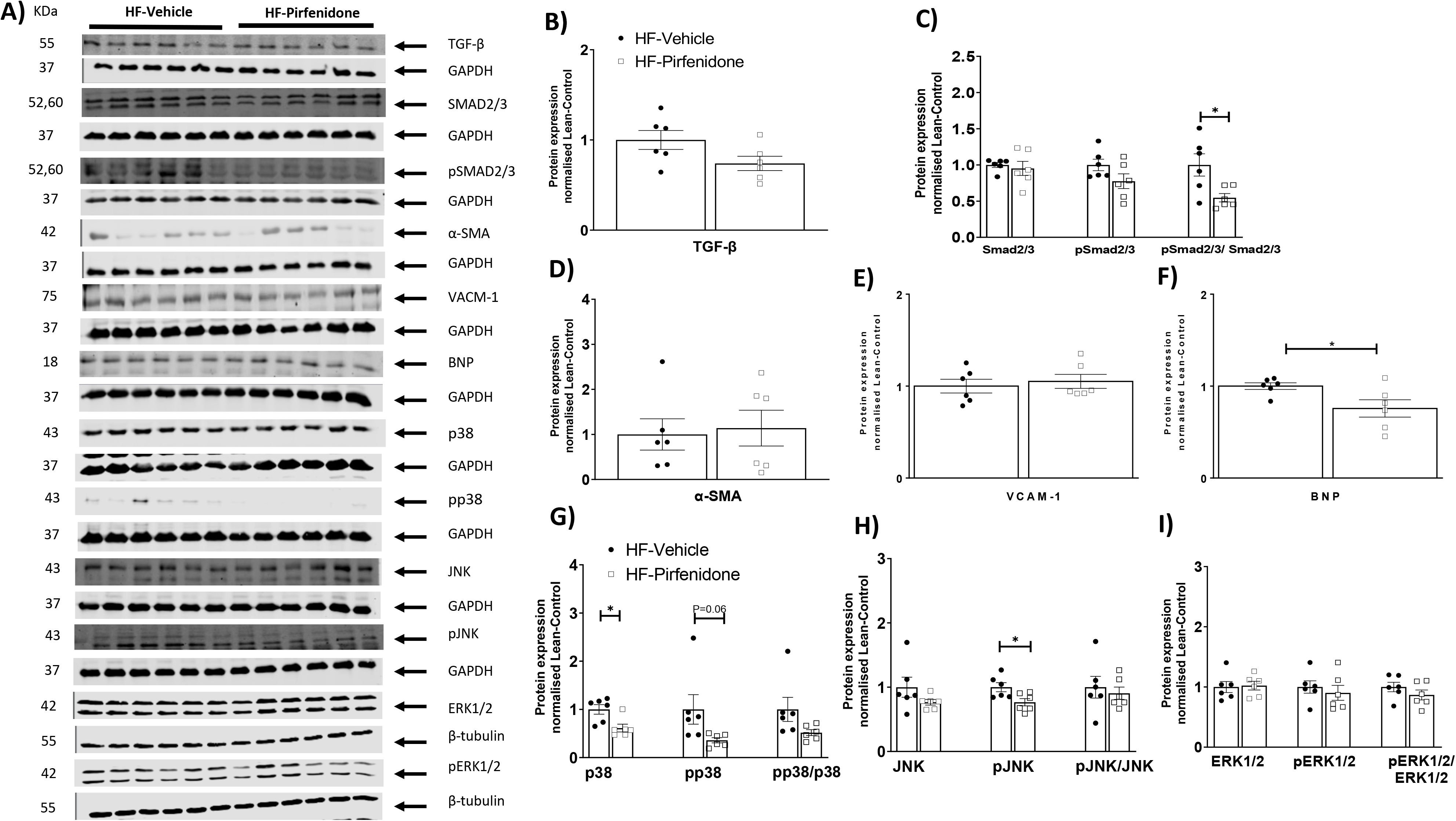
Pirfenidone reduced SMAD2/3 activation, brain natriuretic peptide (BNP) expression, and MAPK activation in the left ventricle of obese mice. C57BL/6 mice were fed with a 60% high fat (HF) diet for 16 weeks. After 12 weeks of HF feeding, mice received twice-daily treatments of vehicle or pirfenidone for 21 days. Protein expression was determined by Western blotting. Representative blots were shown. N=6. \**p*<0.05.

**Figure 8.**
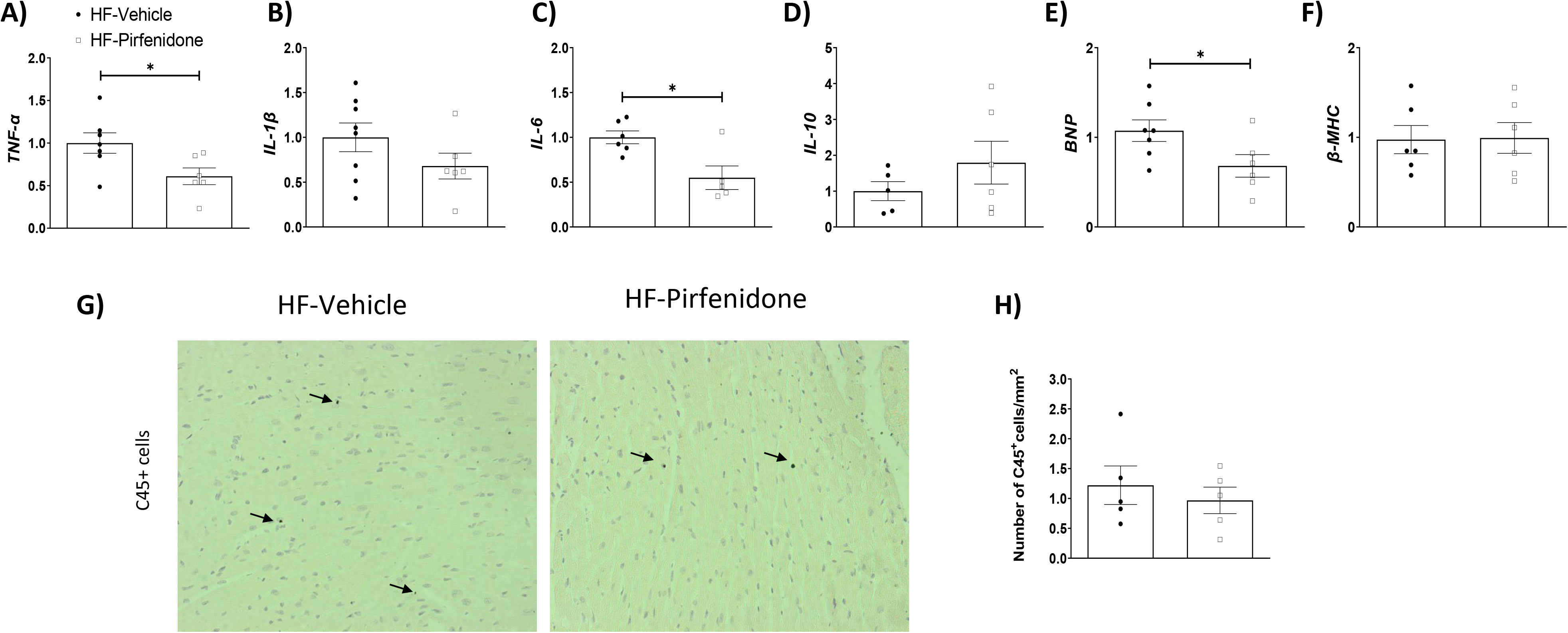
Pirfenidone decreased mRNA levels of pro-inflammatory markers TNF-α and IL-6, and a marker of cardiac function brain natriuretic peptide (BNP) in the left ventricle of obese mice. C57BL/6 mice were fed with a 60% high fat (HF) diet for 16 weeks. After 12 weeks of HF feeding, mice received twice-daily treatments of vehicle or pirfenidone for 21 days. (A-F) mRNA expression was determined by qRT-PCR. N=5-8. (G-H) CD45 positive cells were determined by CD45 immunohistochemistry and quantified by ImageJ. Representative images were shown at 200x magnification (3.10 pixels/μm). N=5. \**p*<0.05. β-MHC: myosin heavy chain β.

## Discussion

Increased ECM deposition has been shown to contribute to obesity-associated insulin resistance in adipose tissue [22], skeletal muscle [4, 5], and liver [6]. The present study demonstrates for the first time a tight association between collagen and hyaluronan deposition in the heart, cardiac insulin resistance, and associated abnormal myocardial remodelling in obese mice. We employed both genetic and pharmacological approaches to modulate ECM deposition in the heart of obese mice. By using these methodologies, we find that increased ECM collagens and hyaluronan in the heart lead to cardiac insulin resistance and cardiac dysfunction while reduction of this increase in ECM components ameliorates cardiac insulin resistance and preserves left ventricular performance in diet-induced obese mice. Glucose is a fuel for the heart particularly during stresses such as ischemia, increased workload, and hypertrophy due to pressure overload. The ability of insulin to promote glucose utilisation in cardiomyocytes directly impacts on cardiac function [2]. Our findings reveal the important niche extracellular components have in heart function. This study outlines a common sequalae that links cardiac fibrosis, insulin resistance, and dynamics. The results herein suggest a simplified therapeutic strategy whereby preventing the dysregulation of the cardiac ECM in obesity and pre-diabetes arrests the progression into cardiac dysfunction.

We show a tight association between increased ECM deposition in the heart and cardiac insulin resistance in obesity, where increased collagen deposition by genetic deletion of MMP9 exacerbates cardiac insulin resistance and decreased hyaluronan deposition by PEGPH20 treatment improves cardiac insulin resistance in obese mice. Using the hyperinsulinaemic-euglycaemic clamp, we measured insulin sensitivity in the heart of conscious mice *in vivo*. This technique overcomes drawbacks of other techniques such as glucose and insulin tolerance tests, which give measures of whole-body metabolism under non-steady state conditions [23]. Others have used *in vivo* PET-CT imaging to measure glucose uptake in the heart, yet these were not insulin-stimulated [24]. Our results are consistent with other studies that showed cardiac remodelling and interstitial fibrosis in obese rodents [25–27]. Here we show that modulation of ECM constitutes in the heart associates with myocardial responses to insulin. This is important as cardiac insulin resistance to glucose utilisation in cardiomyocytes contributes to cardiac dysfunction and the development of heart failure [1, 28]. In a model of abdominal aortic constriction-induced cardiac hypertrophy where cardiomyocytes are enlarged, cardiac-specific insulin resistance is associated with left ventricular systolic and diastolic dysfunction, even in the absence of systemic insulin resistance [29].

The relationship between increased ECM deposition and cardiac insulin resistance extends to cardiac dysfunction. Using PV loop analyses, we show that in the setting of HF diet feeding, PV loops become narrower and taller (Supplemental Fig 3), suggesting that the heart works with smaller volume (Ved, SV albeit insignificant) and higher pressures (Pes) because of increased afterload (Ea) and potentially preload (Ped). The heart adapts to the lower volume by increasing contractility (dP/dt, ESPVR) to maintain ejection fraction and cardiac output. The enhanced contractility could be a result of increased stiffness due to increased ECM deposition and fibrosis, although diastolic function appeared to be unaffected (Tau, EDPVR). PEGPH20 treatment in HF-fed mice preserved both systolic and diastolic pressures and volumes, suggesting that PEGPH20 prevented changes in preload and afterload without the need to adapt for the maintenance of cardiac output and stroke volume. Previous studies using similar diet, 60% HF diet over 16 weeks, showed time-dependent increases in Tau, EDPVR and Ped, indicating diastolic dysfunction [30]. However, in our hands 16 weeks of HF diet did not initiate the same degree of cardiac changes. In the study by Tong et al. [30], mice that did not gain 20% of body weight after 2 months of HF diet consumption were excluded from the analysis, whereas we included all experimental mice in our study. Therefore, we believe our data more accurately represent the heterogeneity of mouse’s response to HF diet, enhancing its translatability to humans. Regardless, our data indicate early stages of cardiac dysfunction, with increased duration of HF diet feeding which may lead to diastolic dysfunction and heart failure.

Hyaluronan, a polysaccharide constituent of the ECM is associated with cardiac ECM remodelling and has been implicated in the cardiac defects of obesity [31]. Increased hyaluronan in the heart of hyaluronidase 2-deficient mice leads to cardiac fibrosis and impaired diastolic function [32]. Likewise, disruption of hyaluronan catabolism causes cardiac abnormalities in patients with a hyaluronidase 2 mutation [33]. Pharmacological removal of hyaluronan by PEGPH20 has been previously shown to improve muscle insulin resistance and reduce adipose tissue inflammation in obese mice [5]. Together with our findings in the heart, where it is shown to prevent HF diet-induced cardiac hypertrophy, fibrosis, and inflammation, we propose that hyaluronan is a promising target for obesity-related metabolic complications. CD44 and RHAMM are main cell surface receptors of hyaluronan that have been implicated in metabolic regulation and obesity [13, 34, 35]. The role of CD44 and RHAMM in regulating cardiac function and insulin resistance merit further investigation.

In addition to hyaluronan, increased collagen deposition is a characteristic of the hearts of obese individuals [36]. Pirfenidone is one of the two approved anti-fibrotic therapies for idiopathic pulmonary fibrosis, exerting its action through inhibiting collagen expression [18]. Although pirfenidone has been shown to reduce cardiac fibrosis and improve left ventricular function in pre-clinical models of myocardial infarction [20], its effect under obese condition had not been studied. Herein, we show that pirfenidone ameliorates cardiac as well as systemic insulin resistance in obese mice, which may contribute to its beneficial effects in cardiac function. A recent clinical trial has shown that pirfenidone reduces myocardial extracellular volumes in patients of heart failure with preserved ejection fraction [37], suggesting a high potential of repurposing pirfenidone for this condition. Our results provide further insight into the beneficial effects of pirfenidone on cardiac insulin resistance.

There is clinical evidence supporting our novel concept that ECM components may specifically drive metabolic and cardiac dysfunction in patients with cardiovascular conditions. Midwall fibrosis is an independent predictor of mortality in patients with moderate and severe aortic stenosis [38]. Excessive myocardial collagen cross-linking determined by the ratio of insoluble and soluble collagen is associated with hospitalisation for heart failure or cardiovascular death in patients with heart failure and arterial hypertension [39]. Moreover, a recent single-cell transcriptomic analysis reveals that fibroblast subtype changes and ECM remodelling highly correlate to disease progression at late stage of pathological cardiac hypertrophy [40]. Although these studies did not specifically examine patients with obesity, myocardial fibrosis and left ventricular hypertrophy are common features of subjects with abdominal obesity [41]. Therefore, myocardial fibrosis may provide a structural basis for pathological changes in the heart and ultimately account for the appearance of adverse cardiovascular events and outcomes [42].

In conclusion, our study establishes a novel link between increased ECM deposition, cardiac insulin resistance and cardiac dysfunction in obesity. By using two anti-fibrotic agents, which target two distinct ECM components, collagen and hyaluronan, our results show that reduction of ECM excess is sufficient to ameliorate cardiac insulin resistance and associated functional changes. These results highlight that early cardiac ECM remodelling in obesity-associated cardiac dysfunction can be mitigated and reversed. We propose that an intervention that prevents deleterious ECM expansion may be protective from further progression to severe cardiovascular consequences.

## Acknowledgement

PEGPH20 is an in-kind gift from Halozyme Therapeutics, Inc under a Material Transfer Agreement.

## Author contribution

VM and LK contributed to the research concept, experimental design, data collection, analysis, and interpretation, and wrote the manuscript. CM, AB, AH, and CH contributed to experimental design, data collection, analysis, and interpretation, and reviewed and edited the manuscript. BD, CC and DW contributed to discussion, and reviewed and edited the manuscript.

---

**Supplemental Table 1.**
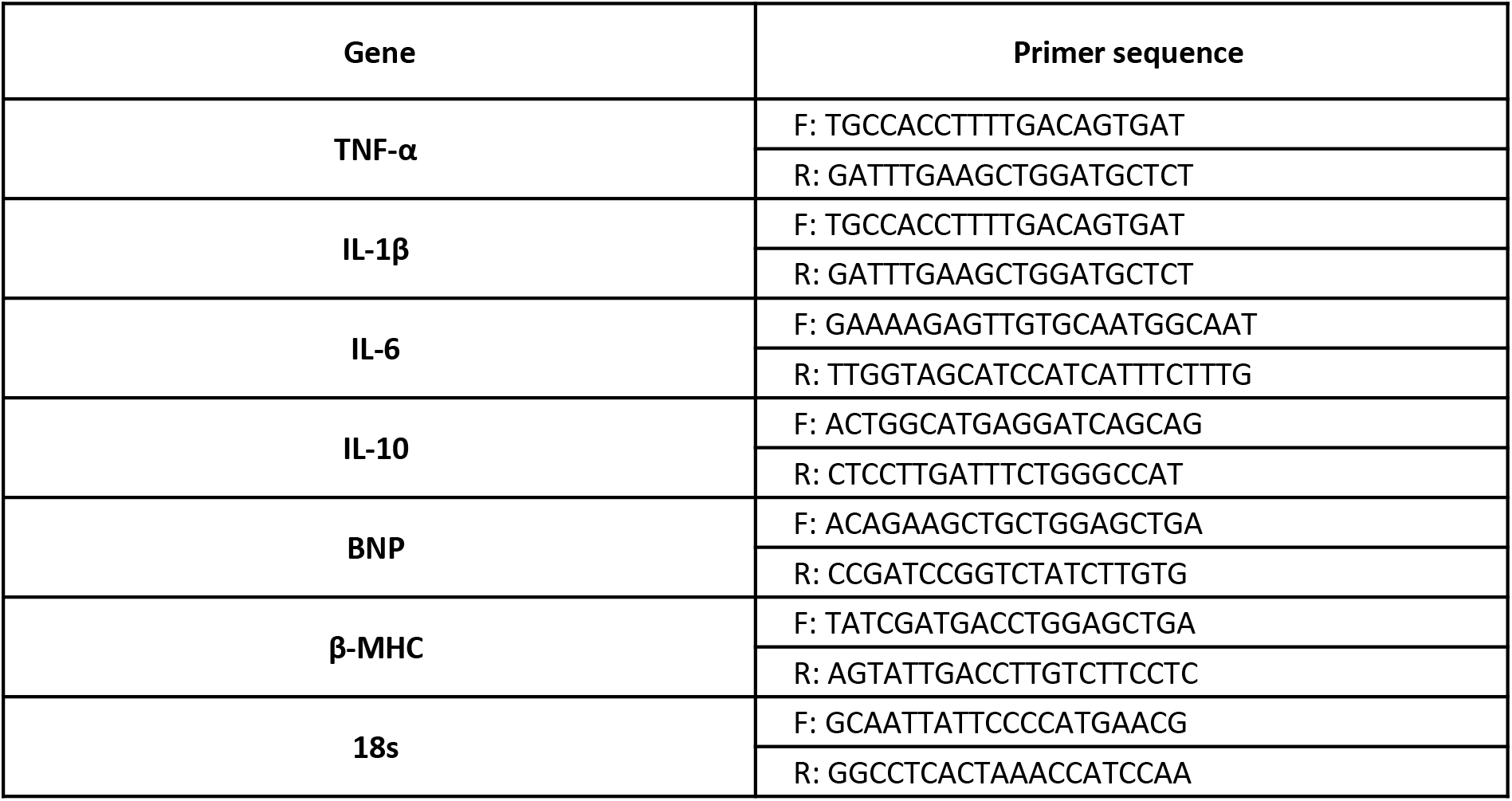
mRNA expression was determined by qRT-PCR using primers shown in the table.

**Supplemental Figure 1.**
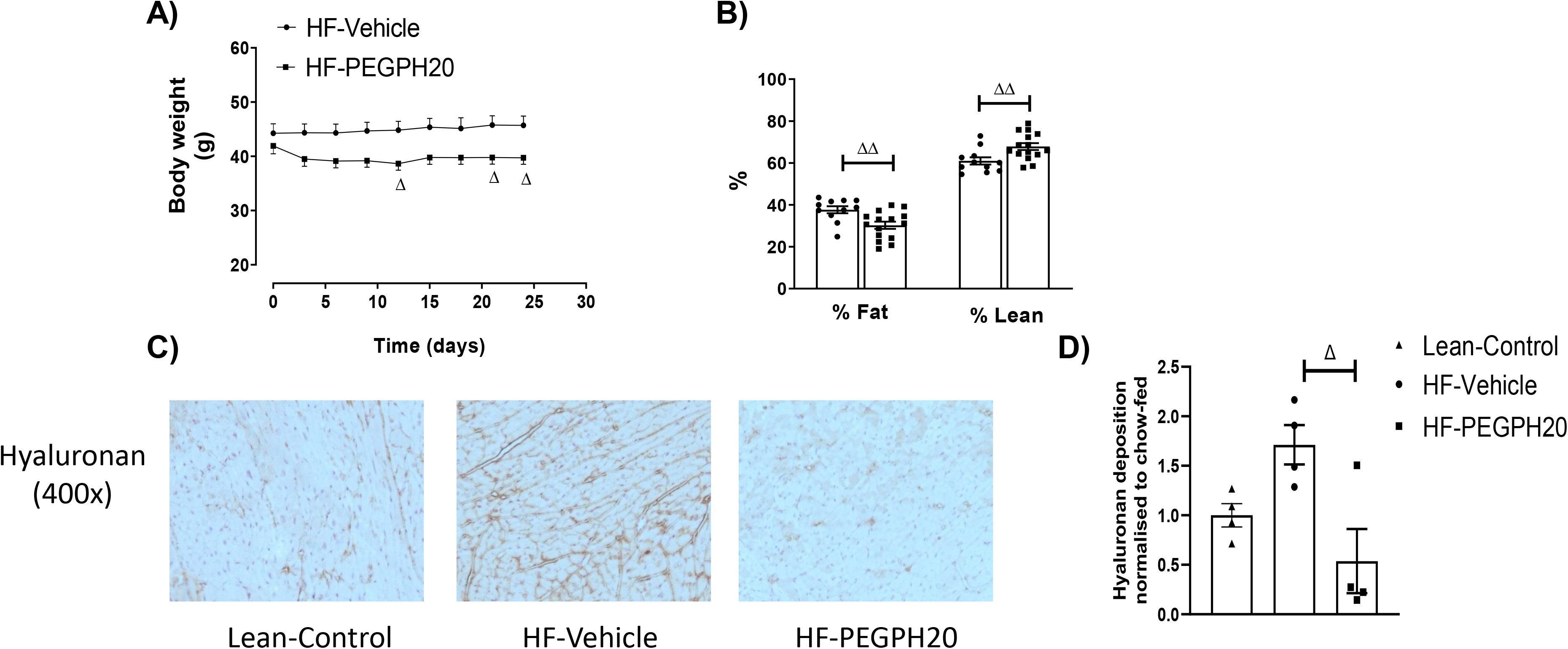
PEGPH20 treatment decreased body weight and %fat mass but increased %lean mass in high fat (HF) diet-fed mice. C57BL/6 mice were fed a 60% HF diet for 16 weeks. After 12 weeks of feeding, mice received either vehicle or PEGPH20, once every 3 days for 24 days. (A) Body weight was monitored daily. N=11-15. (B) Body composition was determined after the vehicle/drug treatment. N=11-15. (C) Hyaluronan was detected by immunohistochemistry and quantified by ImageJ in left ventricle sections. Representative images were shown at 400x magnification (6.21 pixels/μm). N=4. ^Δ^*p*<0.05, ^ΔΔ^*p*<0.01 compared with HF-Vehicle.

**Supplemental Figure 2.**
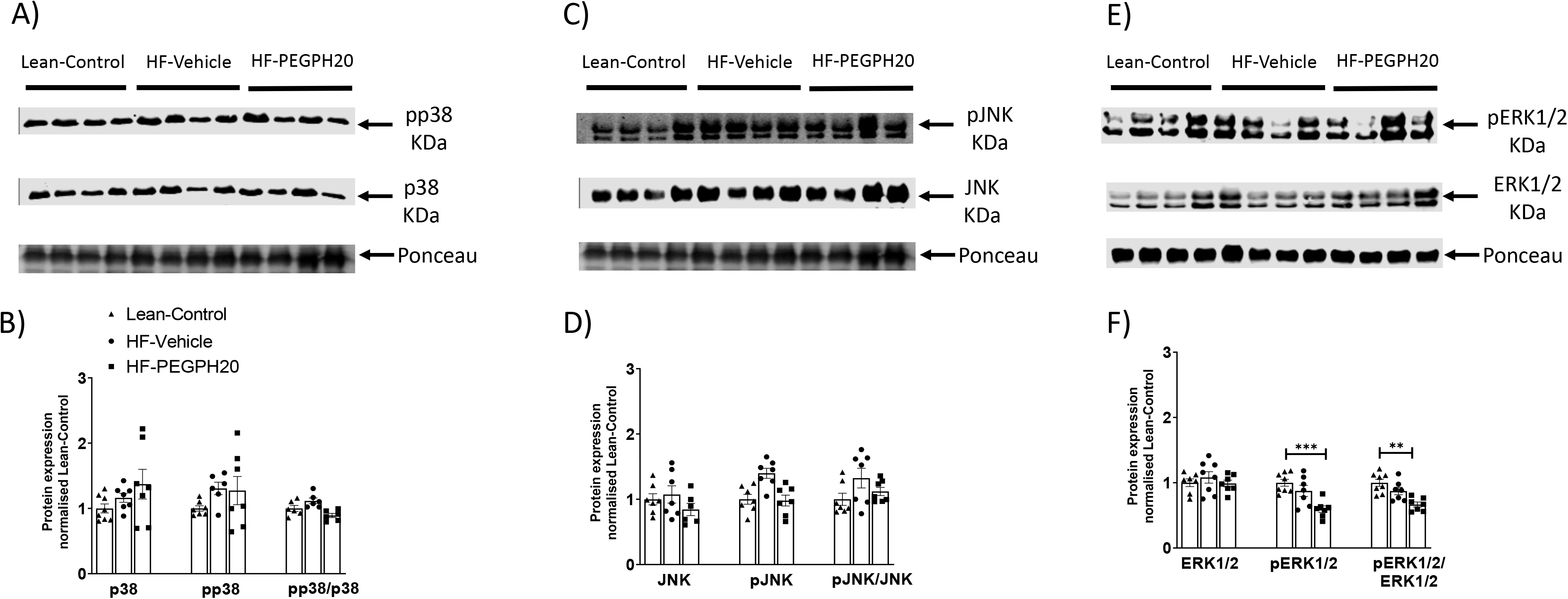
MAPK signalling was not altered by high fat (HF) feeding or PEGPH20 treatment, except that pERK1/2 and ratio of pERK1/2 and total ERK1/2 were lower in HF-PEGPH20 mice relative to lean control mice. Protein expression was determined by Western blotting. Representative blots were shown. N=6-8. \*\**p*<0.01, \*\*\**p*<0.005.

**Supplemental Figure 3.**
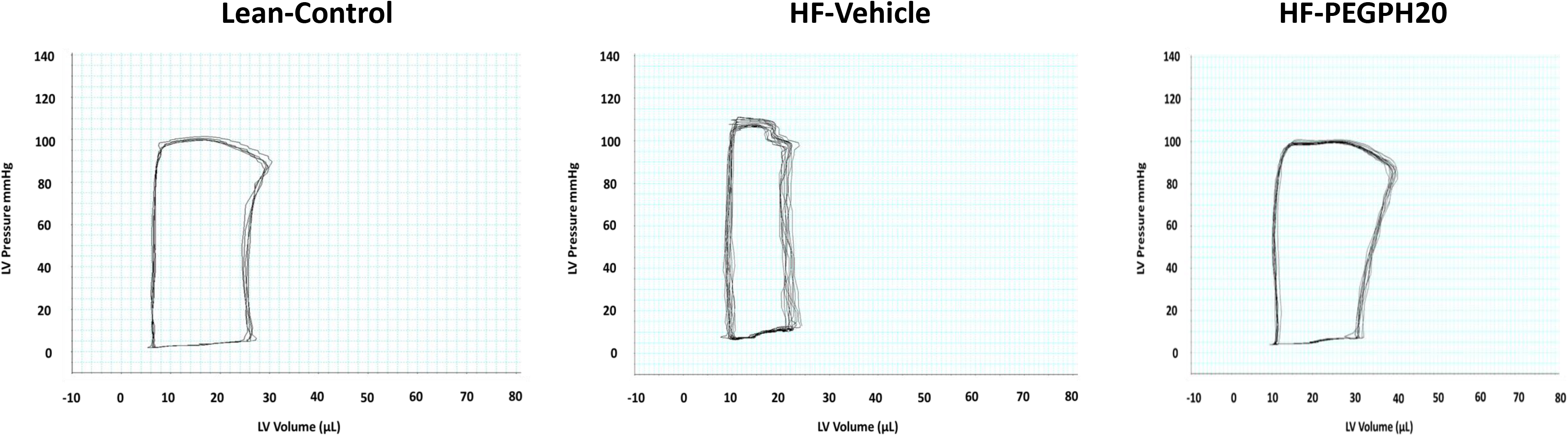
Representative pressure-volume loops. HF-Vehicle mice had taller and narrower loops, showing higher pressures (Pes) and lower volumes (Ved and SV). PEGPH20 treatment reversed these cardiac changes.

